# Sex-specific insights into drug-induced lifespan extension and weight loss in mice

**DOI:** 10.1101/2024.11.30.626149

**Authors:** Aleksey V. Belikov, Angelo Talay, João Pedro de Magalhães

**Affiliations:** Genomics of Ageing and Rejuvenation Lab, Institute of Inflammation and Ageing, University of Birmingham, Birmingham, UK

**Keywords:** Lifespan-extending compounds, longevity drugs, anti-ageing compounds, geroprotectors

## Abstract

The DrugAge database serves as a comprehensive resource for the study of compounds that increase lifespan in model organisms. In the latest version of DrugAge, we have implemented multiple updates, predominantly focusing on mouse (*Mus musculus*) studies to enhance data accuracy and consistency. Key improvements include the re-recording of mouse data from original sources, standardization of drug dosages to parts per million (ppm), and detailed recording of administration routes, treatment initiation ages, and durations. The user interface has also been upgraded. Additionally, weight change data were included to address the potential impact of caloric restriction induced by drug administration on lifespan. Our analysis revealed significant correlations between weight loss and lifespan extension in male mice, particularly in studies conducted by the Interventions Testing Program (ITP), highlighting the importance of considering weight change in lifespan studies. We also observed notable sex-related differences in lifespan and weight change responses, underscoring the need for gender-specific analyses in aging research.

## Introduction

DrugAge (https://genomics.senescence.info/drugs/) is a comprehensive database that compiles a wide range of approved and investigational drugs, chemical compounds, and plant extracts, collectively referred to as “compounds,” which have shown effects on the lifespan of healthy model organisms such as mice (*Mus musculus*), flies (*Drosophila melanogaster*), and worms (*Caenorhabditis elegans*)^1,2^. The database records both significant and non-significant extensions of mean, median, or maximal lifespan. Maintained with rigorous standards, DrugAge ensures all entries are manually curated from the scientific literature. The focus is on compounds potentially impacting aging, and we excluded those tested on disease-prone animals or under harmful conditions. Most negative results are excluded unless particularly relevant to the field of aging research. Each entry corresponds to a specific observation from a study, allowing for multiple, potentially conflicting entries for a single compound, thus encouraging users to interpret the data independently.

The DrugAge homepage facilitates direct searches for specific compounds, species, or bibliography references and allows database downloads. The Browse page displays all entries, which can be filtered or sorted by criteria such as compound, species, strain, dosage, gender, and lifespan changes. PubMed references are provided for further exploration. Individual compound and species pages offer a summary table listing lifespan changes across experiments, and a results table showing detailed data. DrugAge provides visual data representation through lifespan charts, which can toggle between average and maximal lifespan changes, displaying significant and non-significant results or distinguishing male versus female results. The Drug Data Summary page features pie charts illustrating compound and species representations across studies, while the Statistics page offers a summary of assays, compounds, species, references, and the maximum lifespan changes observed for each species. DrugAge Build 5 features over 1000 compounds evaluated in over 3400 experiments across 35 species, supported by more than 660 references.

One of the major updates to Build 5 was the inclusion of weight change data. Caloric restriction is the most reproducible longevity intervention demonstrating the magnitude of the effect on lifespan higher than any compound tested. Caloric restriction may work by preventing obesity^3^, by retarding growth and development if started early in life^4^ or by other mechanisms^5^. In fact, the magnitude of lifespan extension under caloric restriction has r=0.99 correlation with weight gain in control mice across various strains^6^. Compounds may change the smell, taste or appearance of chow to which they are usually added in murine experiments, leading to reduced food consumption and inadvertent caloric restriction. Moreover, some compounds can affect the appetite or feeding behaviour by interfering with dopamine reward systems, leptin levels or circadian rhythms. In fact, it has been proposed that many of the well-known longevity drugs are caloric restriction mimetics^3,7^. Thus, controlling for weight change while testing the effects of various compounds on lifespan is crucial. Here, we demonstrate that pharmacological lifespan extension significantly correlates with weight loss in male mice, particularly in studies conducted by the Interventions Testing Program (ITP)^8,9^.

### Build 5 updates

For build 5 of DrugAge we implemented multiple updates and improvements. Most of the changes concerned data from murine studies. All mouse data was re-recorded from source papers, both for previously included and new studies, to increase accuracy and consistency. Drug dosages were converted to ppm (parts per million, 1 ppm = 1 mg/kg, or 1 μg/g, or 1 mg/l, or 1 μg/ml, or 0.0001% solution) as per ITP standard^9^. Administration route and dosing description were standardised and clearly indicated as “food”, “water”, “bodyweight”, “injection” or “gavage”, reflecting four major types of drug administration and dosing in rodents – ppm of chow, ppm of drinking water, ppm of bodyweight injection and ppm of bodyweight gavage. Age at initiation and treatment duration were recorded from source papers. Separate statistical significance was recorded for average/median and maximum lifespan extension, where available. Weight change was calculated from tables or approximated from plots where data tables or other exact values were not available. Typically, the timepoint with the maximal weight difference between the drug and the control was chosen. If that was not clear from the publication, then the median survival timepoint was selected. Weight change statistical significance was also recorded where available. ITP studies were clearly marked, as the ITP is considered a gold standard in murine lifespan studies due to the use of three independent experimental sites and large cohorts of genetically heterogeneous mice^8,9^.

In the whole database, including other species, some compounds’ duplicates with different names but the same PubChem ID were merged. This explains slight reduction in the number of drugs compared to the previous build despite adding new experiments. Plant and other extracts were uniformly named to contain “extract” in their name, to avoid duplicates and confusion with isolated chemicals. Gender and significance data were converted from string format to multiple choice (dropdown menu) format to remove duplicates and facilitate data selection.

Regarding the user interface, we added a “Gender” column and an option to indicate male and female data on the graph with distinct colours. Sex differences in lifespan effects are often dramatic so these additions were crucial. Please note that other species beyond mice might not always have gender data. “MEAN(Avg Lifespan Change %)” and “MEAN(Max Lifespan Change %)” columns were added to the Summary tables, in addition to the previously available MAX option. This allows users to rank drugs more confidently without the risk of relying on outliers. We made graphs to display the mean and standard deviation of lifespan changes for each compound, combining individual experiments together, which greatly reduced the length and complexity of the graphs. It is also now possible to select only significant results to be displayed on the graph. Additionally, graphs were made to always display zero on the “Lifespan Change (%)” axis to more consistently reflect the relative magnitude of effects, especially in cases where all drugs are very effective, and the poorly performing comparators are missing. Numerous graph and table display issues were also fixed. For *Mus musculus* page specifically, we added “Age at initiation” (search), “Treatment duration” (search), “Max Lifespan Significance” (dropdown), “Weight Change (%)” (slider), “Weight Change Significance” (dropdown) and “ITP” (dropdown) columns to display the expanded data recorded from source articles.

### Murine studies statistics

DrugAge Build 5 *Mus musculus* section features 134 compounds evaluated in 373 experiments, where one experiment refers to a particular dosage and treatment scheme of a compound tested in a particular gender in a particular strain. DrugAge lists 198 experiments for male mice and 172 for females, and males also have a higher number of experiments with significant average/median lifespan extension – 72 (36%) vs 50 (29%) for females. Top 10 experiments in *Mus musculus* ranked by significant average/median lifespan extension are listed in Table 1 for males and Table 2 for females. As can be seen, most suscessful treatments were initiated within the first 4 months of life, with the exception of rapamycin, X203 (anti-IL-11) and 17-alpha-estradiol, which were initiated at 9 months or later. Most compounds were delivered in chow, as this is the easiest option. Many compounds increasing average/median lifespan also extended maximum lifespan. Most of the top experiments are not from the ITP, with independent validation pending. Moreover, several of them are from 1970s, 1980s, or even 1960s and 1950s, and require reproduction in modern labs. Many compounds induced weight loss, which will be discussed in the next section.

**Table 1.**
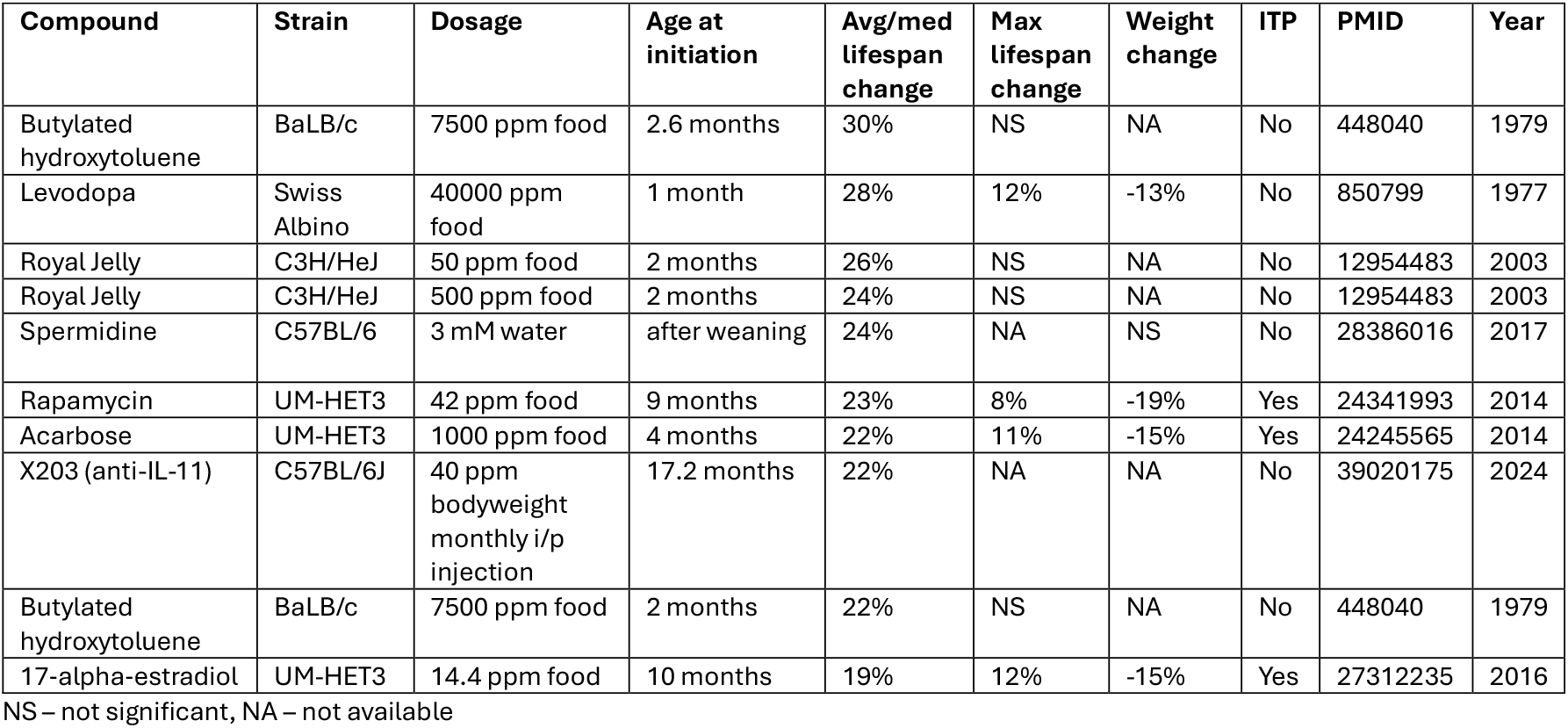
Top 10 experiments in *Mus musculus* ranked by significant average/median lifespan extension in males.

**Table 2.** Top 10 experiments in *Mus musculus* ranked by significant average/median lifespan extension in females.

| Compound | Strain | Dosage | Age at initiation | Avg/med lifespan change | Max lifespan change | Weight change | ITP | PMID | Year |
| --- | --- | --- | --- | --- | --- | --- | --- | --- | --- |
| Metformin | SHR | 100 ppm water | 3 months | 38% | 21% | NS | No | 18728386 | 2008 |
| Pineal gland extract | C3H/Sn | 0.5 mg subcutaneous injection 5 days monthly | 3.5 months | 31% | 14% | NA | No | 6752596 | 1982 |
| Thymus extract | C3H/Sn | 0.5 mg subcutaneous injection 5 days monthly | 3.5 months | 28% | 11% | NA | No | 6752596 | 1982 |
| 2-mercaptoethylamine hydrochloride | C3H | 10000 ppm food | after weaning | 26% | NA | -1% | No | 13711616 | 1961 |
| Rapamycin | UM-HET3 | 42 ppm food | 9 months | 26% | 11% | -20% | Yes | 24341993 | 2014 |
| X203 (anti-IL-11) | C57BL/6J | 40 ppm bodyweight monthly intraperitoneal injection | 17.2 months | 25% | NA | NA | No | 39020175 | 2024 |
| Butylated hydroxytoluene | BaLB/c | 7500 ppm food | 2.6 months | 25% | NS | NA | No | 448040 | 1979 |
| Rapamycin | UM-HET3 | 14 ppm food | 9 months | 21% | 11% | -4% | Yes | 24341993 | 2014 |
| Phenformin | C3H/Sn | 2 mg oral gavage 5 times a week | 3.5 months | 21% | 28% | NA | No | 14618027 | 2003 |
| Calcium pantothenate | C57BL/6 | 300 µg/day in water | 1 month | 20% | NA | 12% | No | 13614445 | 1958 |

NS – not significant, NA – not available

Effects of DrugAge-listed compounds on the average/median murine lifespan range from -30 to +41 percent of control, resulting in 4.4% extension on average (Fig 1A). Effects on the maximum lifespan range from -18 to +31 percent of control, resulting in 2.6% extension on average (Fig 1A). When only statistically significant effects on the average/median murine lifespan are considered, they range from -17 to +38 percent of control, resulting in 10.7% extension on average (Fig 1B). Significant effects on the maximum lifespan range from -14 to +28 percent of control, resulting in 10% extension on average (Fig 1B). This might reflect the size of the effect (around 10%) required to achieve statistical significance in a typical murine lifespan experiment.

**Figure 1.**
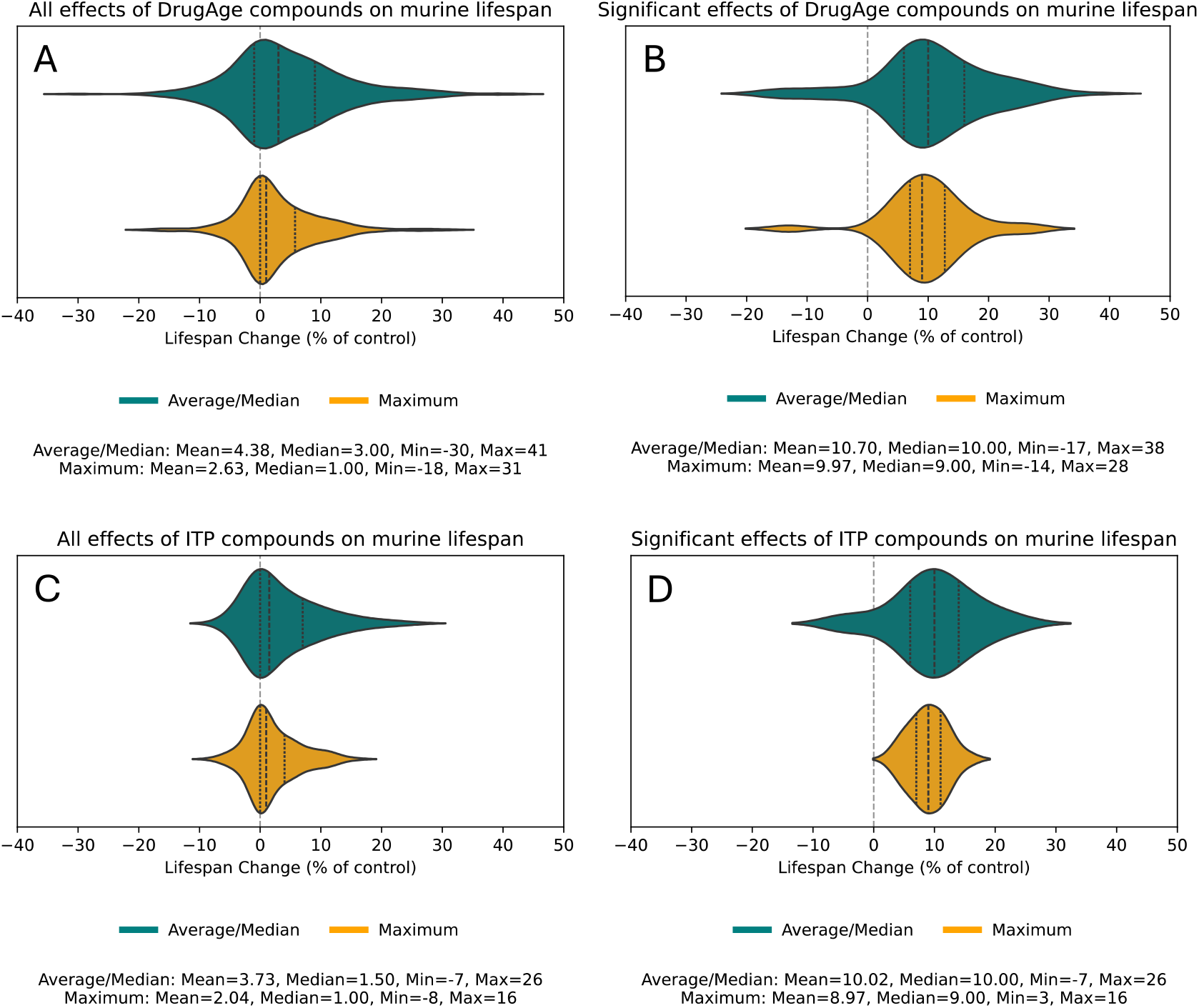
Effects of compounds on average/median and maximum murine lifespan. (**A, B**) Compounds from DrugAge. (**C, D**) Compounds from ITP studies. (**A, C**) All effects. (**B, D**) Significant effects.

When only the ITP studies are considered, effects on the average/median murine lifespan range from -7 to +26 percent of control, resulting in 3.7% extension on average (Fig 1C). Effects on the maximum lifespan in ITP studies range from -8 to +16 percent of control, resulting in 2% extension on average (Fig 1C). Less negative effects on the lifespan in ITP studies are likely explained by carefull selection of compounds for testing, usually based on previous successful studies^8,9^. Finally, when only statistically significant effects on the average/median murine lifespan in the ITP studies are considered, they range from -7 to +26 percent of control, resulting in 10% extension on average (Fig 1D). Significant effects on the maximum lifespan in ITP studies range from +3 to +16 percent of control, resulting in 9% extension on average (Fig 1D).

The average/median lifespan change positively correlates with the maximum lifespan change, both in males and females, with females consistently achieving higher and more significant correlation (Supplementary Fig 1). The correlation is very strong in the full ITP dataset, both in males (slope: 0.46, R^2^: 0.47, p-value: 7×10^−13^) and females (slope: 0.58, R^2^: 0.57, p-value: 5×10^−16^) (Supplementary Fig 1C). This might be due to a consistent way of measuring both median (50% survival) and maximum (10% survival) lifespan in ITP studies^8,9^.

### Weight change analysis

When all data are considered, correlations between weight change and median or maximum lifespan change are significant for males and females combined and for males alone but not for females alone (Supplementary Fig 2). Nevertheless, the R^2^ and slopes are small, even for males. However, when ITP-only data are used, p-values, R^2^ and slopes are much larger (Fig 2). In males there is a strong correlation between the increase in lifespan and decrease in weight, both for median (slope: -0.76, R^2^: 0.52, p-value: 3.09×10^−11^, Fig 2C) and for maximum (slope: -0.97, R^2^: 0.35, p-value: 3.73×10^−7^, Fig2D) lifespan. Curiously, in females many compounds led to dramatic decrease in weight without substantial increase in lifespan (Fig 2E). Correlations might be more pronounced in ITP studies due to higher quality and uniformity of the data, being collected for all drugs at the same sites, in the same conditions, using the same protocols (incl. for weight measurements) and using large cohorts of genetically heterogeneous mice^8,9^.

**Figure 2.**
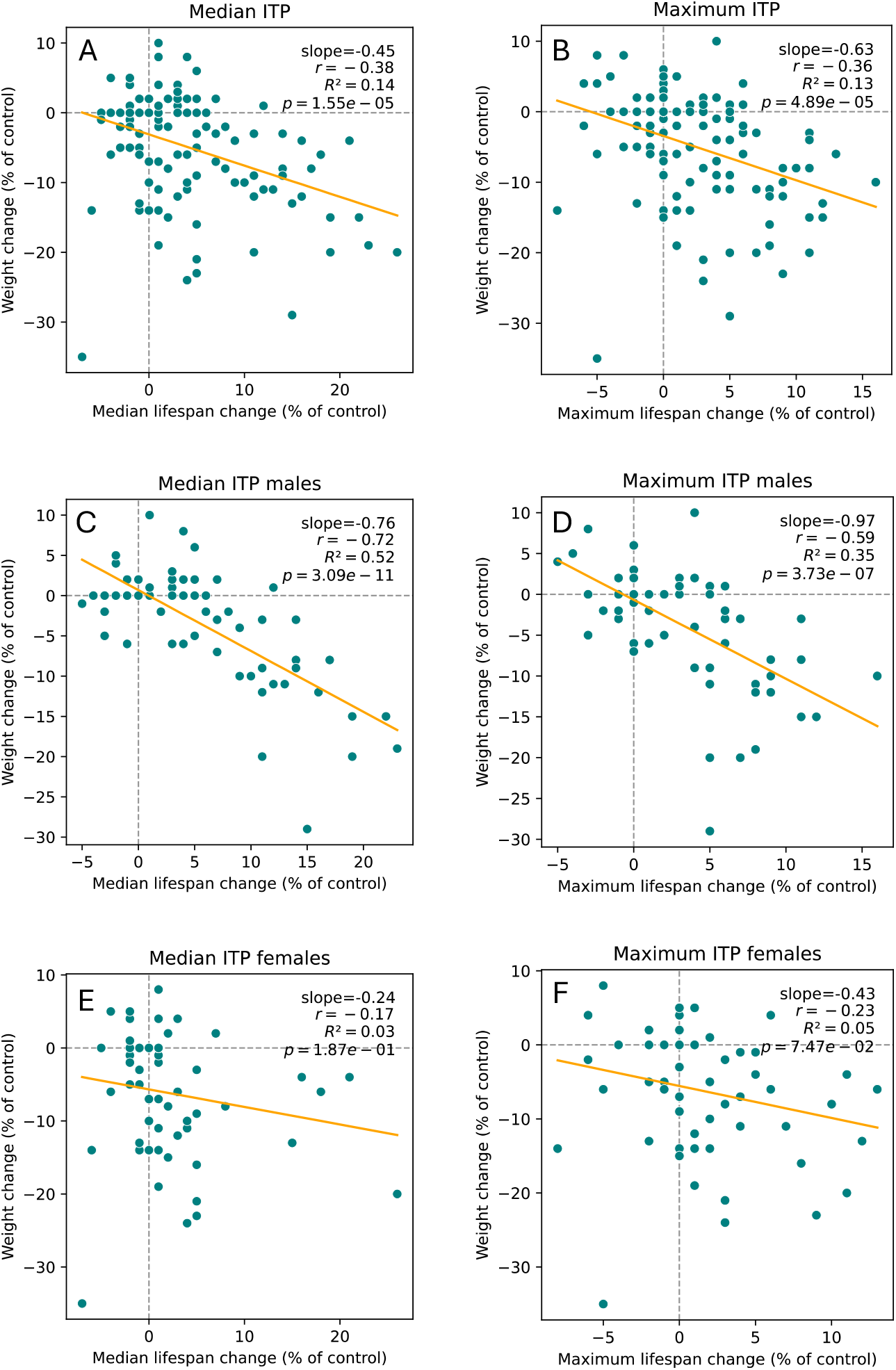
Correlations between weight change and median (**A**,**C**,**E**) or maximum (**B**,**D**,**F**) lifespan change for males (**C**,**D**), females (**E**,**F**) and both sexes combined (**A**,**B**) from ITP studies.

Because we observed larger correlations in males than in females, we decided to look at sex-related differences more closely. We performed pairwise comparisons for each drug that has been tested both in males and females by ITP. When all such compounds are considered, including the ones not extending lifespan significantly, the difference between males and females in median lifespan change (p-value: 4.58×10^−4^, Fig 3A) and weight change (p-value: 1.66×10^−5^, Fig 3C) is more significant than in maximum lifespan change (p-value: 0.0144Fig 3B). For the majority of the compounds, males have higher median lifespan extension than females but less pronounced weight loss, but there are exceptions. Plotting pairwise lifespan and weight changes on the same graph confirms that the same compound can typically induce stronger weight loss in females but stronger median lifespan extension in males (Supplementary Fig 3). The results for maximum lifespan extension are less clear.

**Figure 3.**
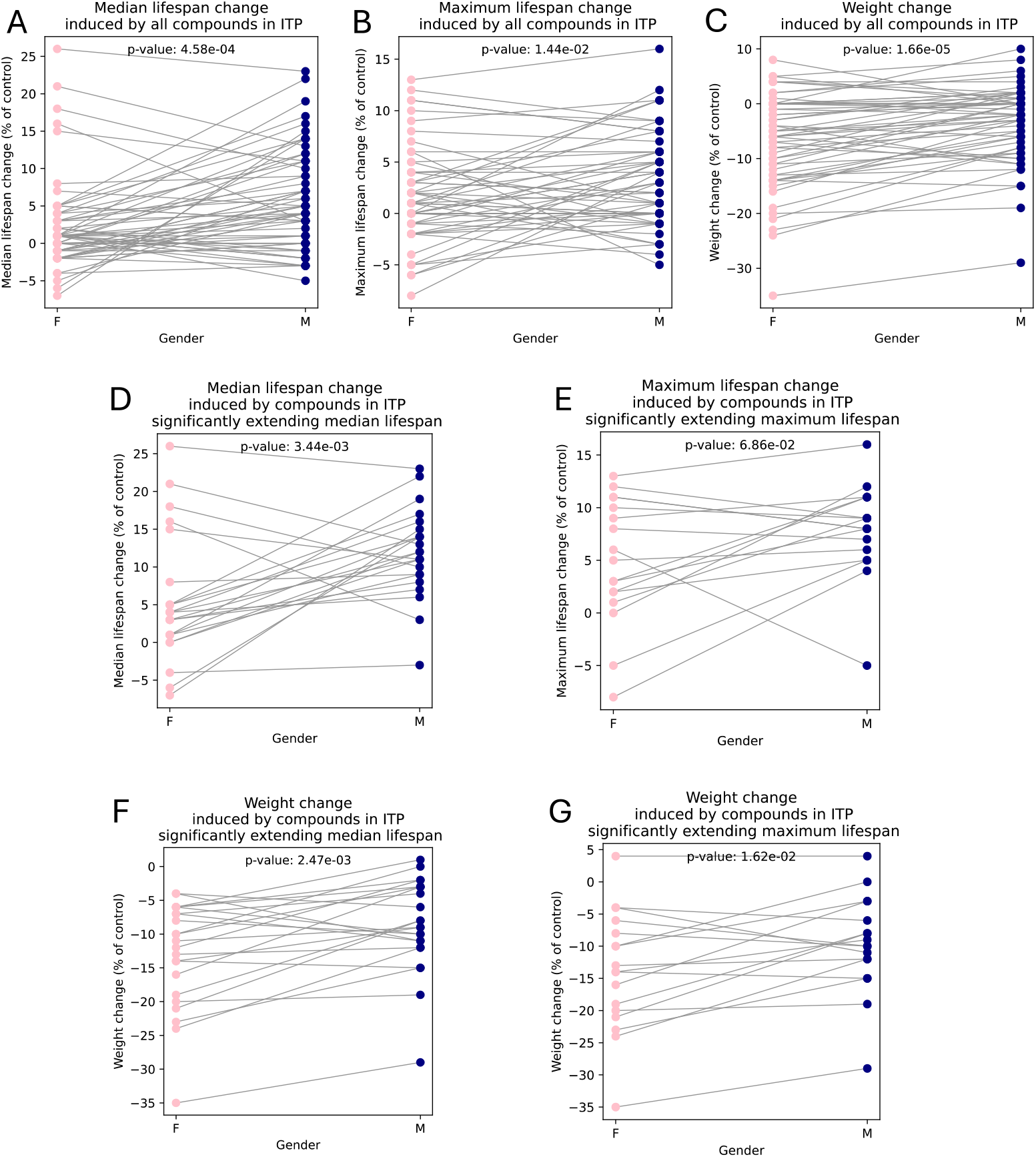
Pairwise comparisons between males (M, *navy*) and females (F, *pink*) of median lifespan changes (**A**,**D**), maximum lifespan changes (**B**,**E**) and weight changes (**C**,**F**,**G**) for compounds from ITP studies. Lines represent individual compounds. (**A**,**B**,**C**) All tested compounds were considered. (**D**,**E**,**F**,**G**) Only compounds significantly extending median (**D**,**F**) or maximum (**E**,**G**) lifespan were considered.

When focusing specifically on compounds significantly extending median lifespan in ITP studies (in at least one gender), the higher median lifespan increase (Fig 3D) and less pronounced weight loss (Fig 3F) in males compared to females remained statistically significant. Curiously, while there was no significant difference between males and females in the magnitude of maximum lifespan increase amongst compounds that significantly extended maximum lifespan in ITP studies (Fig 3E), females still had significantly more pronounced weight loss than males when receiving those compounds (Fig 3G).

## Discussion

Our study examined the relationship between weight change and lifespan extension in mice treated with various compounds, revealing significant sex-specific differences. The results indicate that weight loss correlates strongly with increased lifespan in males but not in females, suggesting that the mechanisms by which drugs affect longevity may differ between sexes.

Several studies in genetically heterogeneous mice (the same kind as used in ITP studies) highlighted a different relationship between bodyweight and lifespan in males compared to females. One study showed that the difference in survival between heavy and light mice is much more pronounced in males than in females, with heavy males on average living less than heavy females, and light males living longer than light females^10^. Another study showed that lifespan negatively correlates with weight in males, especially at younger ages, but there is almost no correlation in females^11^. Notably, that study also clearly demonstrated that survival in the first two thirds of lifespan in much worse for males compared to females^11^. Similarly, in a recent study, long-lived males were lighter than short-lived males, whereas there was almost no difference in weight between long-lived and short-lived females^12^. Consistent with this, in humans, being overweight in adolescence is associated with an increased risk of mortality from all causes, coronary heart disease, atherosclerotic cerebrovascular disease and colorectal cancer among men, but not among women^13^.

Interestingly, male mice have a locus on chromosome 9 which modulates longevity through its effect on growth or body weight, but no such locus has been found in females^12^. Top-scoring genes in this locus are Glb1, which is a beta-galactosidase, and Rtp3, which is a receptor-transporting protein that promotes functional cell surface expression of the bitter taste receptors TAS2R16 and TAS2R43. We speculate that higher expression of Rtp3 in some males thus might make them consume less chow due to more pronounced bitterness and thus reduce weight and promote longevity.

Compounds that decrease weight in animals may function as caloric restriction mimetics^3,7^. The pronounced weight loss without a corresponding increase in lifespan we noted in female mice treated with various compounds suggests that weight reduction is not the main mechanism of lifespan extension for this gender. In the seminal 1935 experiment in rats by McCay, Crowell and Maynard, caloric restriction retarded growth of both males and females but extended lifespan only in males^4^. One alternative to caloric restriction aimed to achieve similar lifespan extension effects is protein restriction, such as restriction of methionine or branched chain amino acids leucine, isoleucine and valine^7^. Lifelong restriction of branched chain amino acids reduced weight in C57BL/6J females more than in males but led to a 30% increase in lifespan and a reduction in frailty only in males^14^. This diet also affected mTOR and FoxO pathways only in males^14^.

This divergence between genders in lifespan responses to compounds, dietary restriction and weight loss may be due to different metabolic, hormonal, developmental or genetic profiles. For instance, females might experience more pronounced metabolic adaptations or hormonal fluctuations in response to caloric restriction that do not necessarily translate to increased longevity. In fact, it has been recently shown in a study of female genetically heterogeneous mice that the top *within-diet* predictor of long lifespan was the ability of mice to *retain* bodyweight under caloric restriction, an indicator of stress resilience^15^. Even more surprisingly, in female mice, lean tissue mass was negatively correlated with lifespan, more strongly early in life, while adiposity was positively correlated with lifespan, more prominently in late life^15^. Moreover, fasting glucose and energy expenditure were not associated with lifespan^15^. This evidence argues against the reduction in obesity or in metabolism as the mechanism for lifespan extension under caloric restriction in female mice^5^. Unfortunately, as males have not been used in that study, it is not clear if these conclusions are unique to females.

Altogether, these findings lend support to the idea that when caloric restriction and its mimetics are able to extend female lifespan, they do so not by decreasing weight but by some other mechanism^16^, however, they almost always work in males, most likely via weight loss and/or growth retardation. This likely explains why most compounds that decrease weight are only able to increase lifespan in males but not in females.

The significant sex-specific differences in median lifespan and weight change observed in our study have important implications for future research. It is crucial to include both male and female subjects in longevity studies to fully understand the effects of various compounds. Future research should aim to elucidate the underlying biological mechanisms that contribute to these differences, such as hormonal influences, metabolic rates, and genetic factors. Additionally, the potential confounding effects of drug-induced changes in feeding behaviour must be carefully controlled and monitored. Compounds that alter the taste, smell, or appearance of food, or that affect appetite and feeding behaviour through neuroendocrine pathways, could inadvertently induce caloric restriction, complicating the interpretation of results. Therefore, controlling for weight change while testing the effects of various compounds on lifespan is essential.

## Conclusions

The updates in Build 5 of the DrugAge database have significantly enhanced the quality and usability of lifespan data, particularly from murine studies. The standardization of drug dosages, detailed recording of treatment protocols, and inclusion of weight change data provide a more robust framework for analysing the effects of compounds on lifespan. The improvements in the user interface, including new data columns and refined graphing options, allow for more precise and comprehensive analyses, facilitating better understanding and comparison of drug effects across different studies. Furthermore, our findings underscore the importance of controlling for weight change, as it can influence the outcomes of lifespan studies, especially in male mice. The gender-specific differences observed in our analysis highlight the necessity of separate evaluations for males and females to interpret the effects of longevity interventions. Overall, these updates make the DrugAge database a more powerful tool for researchers investigating the pharmacological modulation of lifespan and pave the way for more accurate and reproducible findings in the field of aging research.

## Author contributions

AVB compiled data and updated DrugAge, contributed to DrugAge interface improvements, performed analyses, prepared the figures and wrote the manuscript. AT implemented improvements to the DrugAge interface and database. JPM contributed to the design of DrugAge and of the study, interpreted the data and edited the manuscript. All authors read and approved the final manuscript.

