## Supplementary Fig for "Sex-specific insights into drug-induced lifespan extension and weight loss in mice"

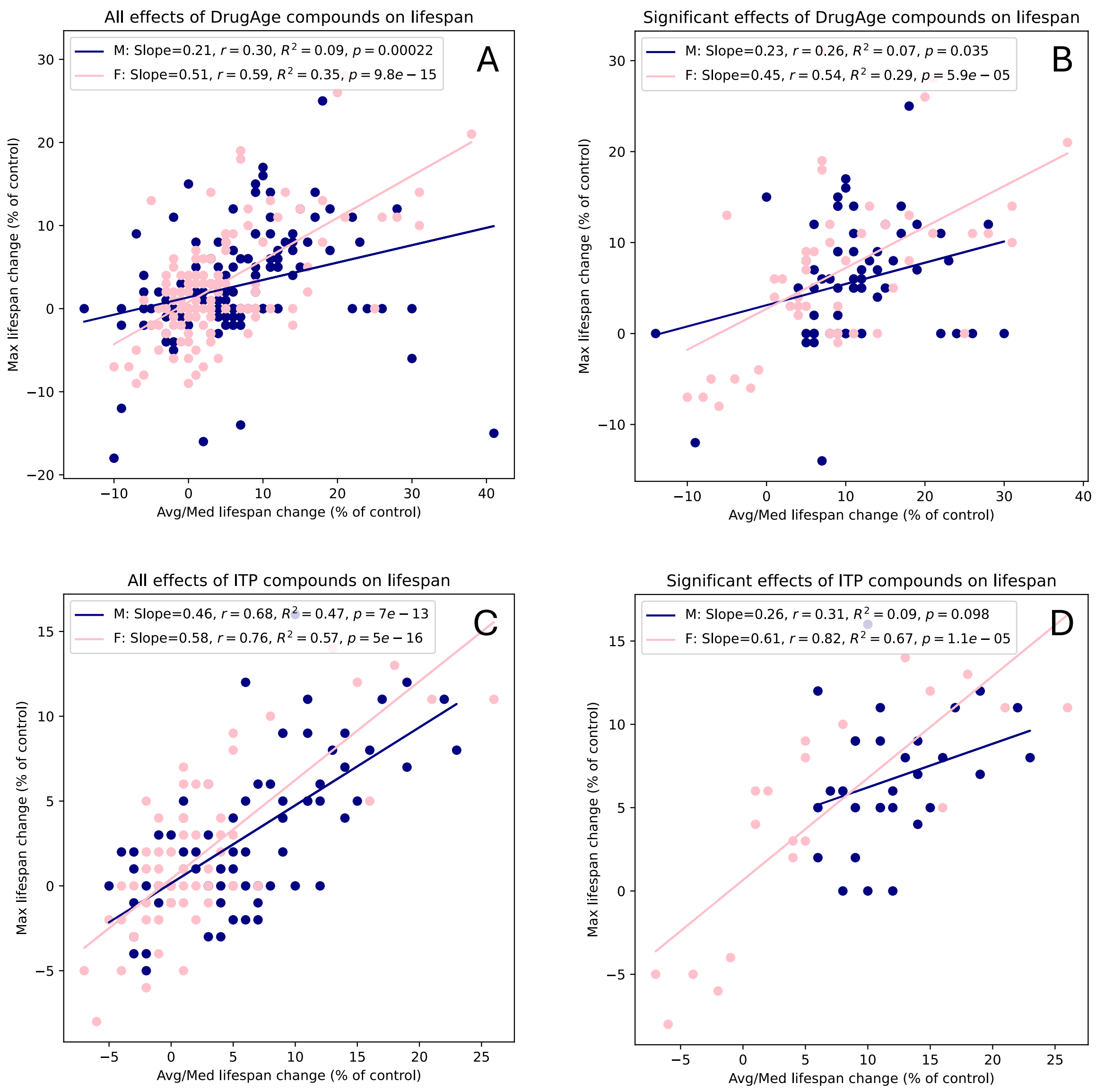


**Supplementary Figure 1.** Effects of compounds on average/median vs maximum murine lifespan for males (M, navy) and females (F, pink). (**A**,**B**) Compounds from DrugAge. (**C**,**D**) Compounds from ITP studies. (**A**,**C**) All effects. (**B**,**D**) Significant effects.

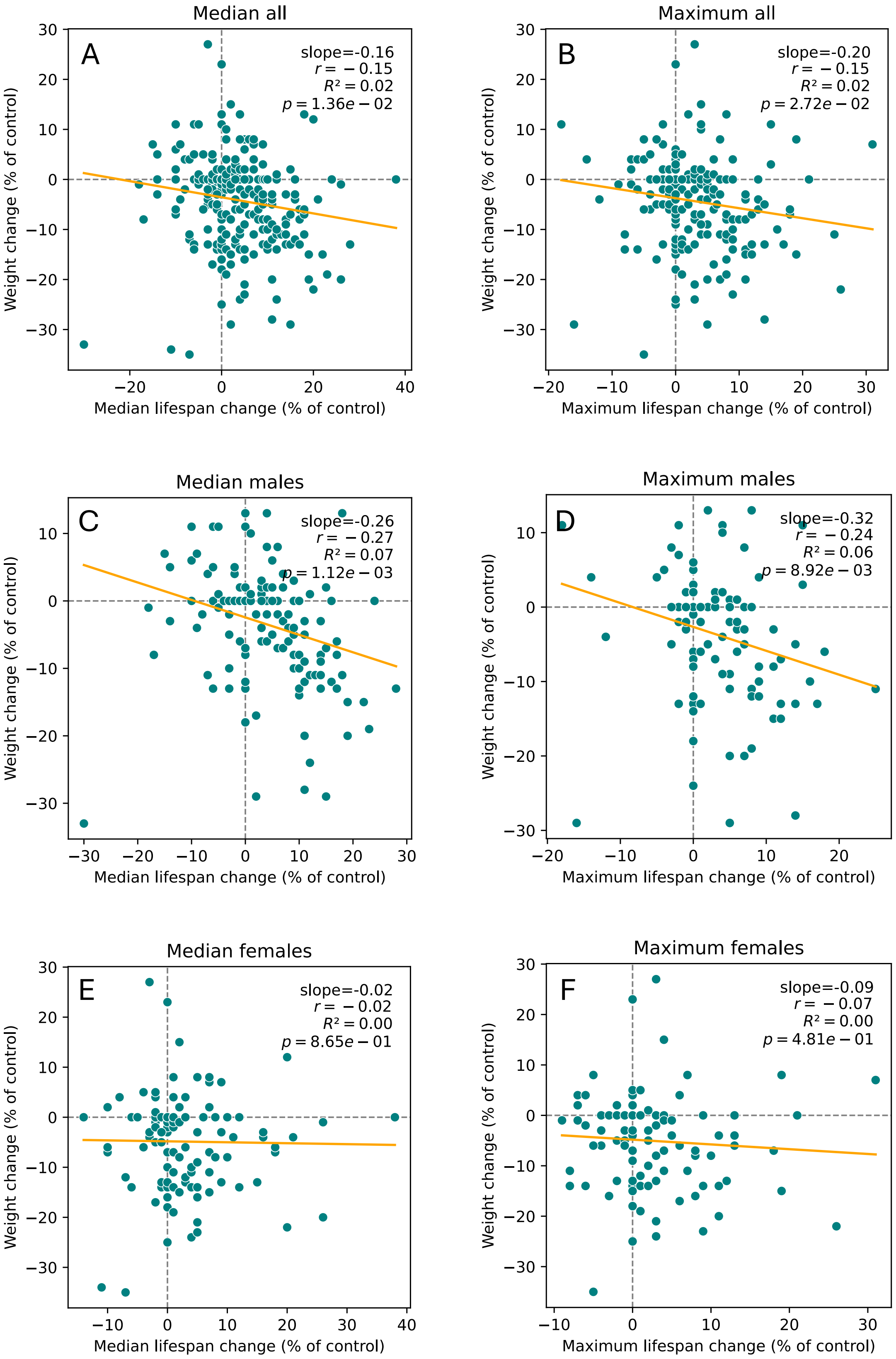


**Supplementary Figure 2**. Correlations between weight change and median (**A**,**C**,**E**) or maximum (**B**,**D**,**F**) lifespan change for males (**C**,**D**), females (**E**,**F**) and both sexes combined (**A**,**B**) from all available murine studies.

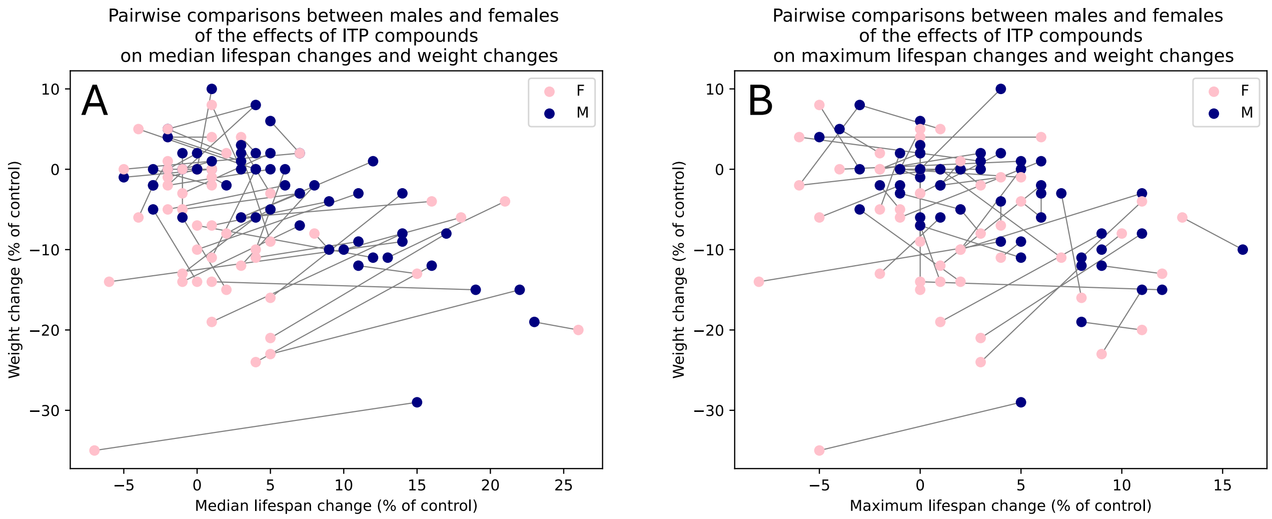


**Supplementary Figure 3.** Pairwise comparisons between males (M, navy) and females (F, pink) of the effects of compounds from ITP studies on median (**A**) or maximum (**B**) lifespan changes and weight changes. Lines represent individual compounds. All tested compounds were considered, including those with non-significant effects.
